# Clinical scoring is the most sensitive biomarker detecting developing sepsis in newborn mice

**DOI:** 10.1101/322776

**Authors:** Beate Fehlhaber, Kathrin Rübensam, Anna S. Heinemann, Sandra Pfeifer, Maren von Köckritz-Blickwede, Dorothee Viemann

## Abstract

Newborn individuals are highly susceptible to infectious diseases. For better insights into age-specific host-pathogen interactions infection models are increasingly employed in neonatal mice. However, for newborn mice no measures are available to objectify the clinical disease state, particularly not in a longitudinal manner, to meet legal animal welfare requirements. We developed a scoring system for newborn mice that relies on observational and examination-based parameters and validated it by applying a *Staphylococcus aureus*-induced infection model in two different mouse strains.

The scoring results strongly correlated with the death kinetics independent of which mouse strain was used. A score above 7 predicted fatality. While the score values increased already at early sepsis stages the large majority of plasma cytokine levels remained comparable to those in uninfected control neonates. The levels of interleukin (Il)-6, chemokine C-C motif ligand 5, Il-1α and tumor necrosis factor α were not increased before 24 hours after infection and correlated only at this late stage of sepsis with the scored disease state.

We propose the first clinical scoring system that serves as important research tool to evaluate the clinical course of sepsis in newborn mice. It detects health impairments of newborn pups in a highly sensitive and longitudinal manner, providing information about the disease severity as well as prognosis.

## Introduction

Neonatal sepsis is still one of the leading causes of death worldwide (1). Despite improved quality control in neonatal care and implementation of bundles for infection prevention, clusters and outbreaks of nosocomial infections in neonatal intensive care units are still a frequent phenomenon (2–4), calling for better understanding of the newborn’s immunity. Moreover, the increasing awareness that the neonatal window represents a crucial period of life for sustainable immunological imprinting (5–12) has initiated numerous investigations focusing on long-term consequences of microbial challenges during early childhood. Better insight into the mechanisms of early life programming and reprogramming of immunity aims at gaining access to novel preventive strategies against the increasing incidence of chronic inflammatory and autoinflammatory diseases (13). To approach to these scientific questions studies in neonatal animals are indispensable.

Evaluating the health status of experimental animals can be realized by using invasive and non-invasive methods. According to European guidelines, the highest priority of keeping, breeding and using animals in academic research is the wellbeing and protection of these animals (14). Therefore, non-invasive methods are preferable to reduce pain and suffering to a minimum (15). For adult mice, a few scoring systems are published, mostly considering specific clinical symptoms of the modeled disease (16–18). Appropriate schemes for the evaluation of the health status and welfare of neonatal rodents are lacking. Moreover, the objectified clinical evaluation of neonatal mice would be helpful to follow individual disease courses, while invasive studies, e.g. blood drawings, are extremely difficult to accomplish in a longitudinal manner. However, scoring systems for adult animals can usually not be transferred to neonatal animals due to the developmental and anatomic peculiarities of newborn pups. In human neonates, clinical evaluation is still more sensitive than laboratory biomarkers to detect septic events (19, 20). Thus, an objective clinical assessment tool for neonatal mice is urgently needed, particularly, when applying models of infection and inflammation.

In this study, we developed a scoring system for neonatal mice that is based on observation and minimal physical examination. The value of the scoring system was studied by applying a model of *Staphylococcus (S.) aureus*-induced neonatal sepsis in two mouse strains with different susceptibility to septic diseases. Evaluation criteria were the correlation between scoring results and disease courses and outcomes, the predictive accuracy of the scoring in terms of fatality and its sensitivity to detect impairments of the health status in newborn mice compared to common invasive laboratory markers of inflammation.

## Results

### Novel score sheet for neonatal mice

Due to obvious developmental differences between adult and neonatal mice we developed an adapted score system for neonates considering the lack of fur, accessibility of skin color, reduced movement, nursing/seeking behavior and visibility of abdominal milk spots. The evaluation of newborn animals included observational and examination-based parameters to assess different categorical aspects of the health status, i.e. pain, appearance, clinical symptoms, spontaneous behavior and provoked behavior. The parameters were listed in a score sheet (Fig. 1) that was used to record the scores awarded to individual mice at defined time points. The score values were summed up to a final total score as described in Materials and Methods. In Fig. 2, healthy and diseased neonates are shown to illustrate exemplarily clear differences in selected observational scoring parameters.

**FIG 1.**
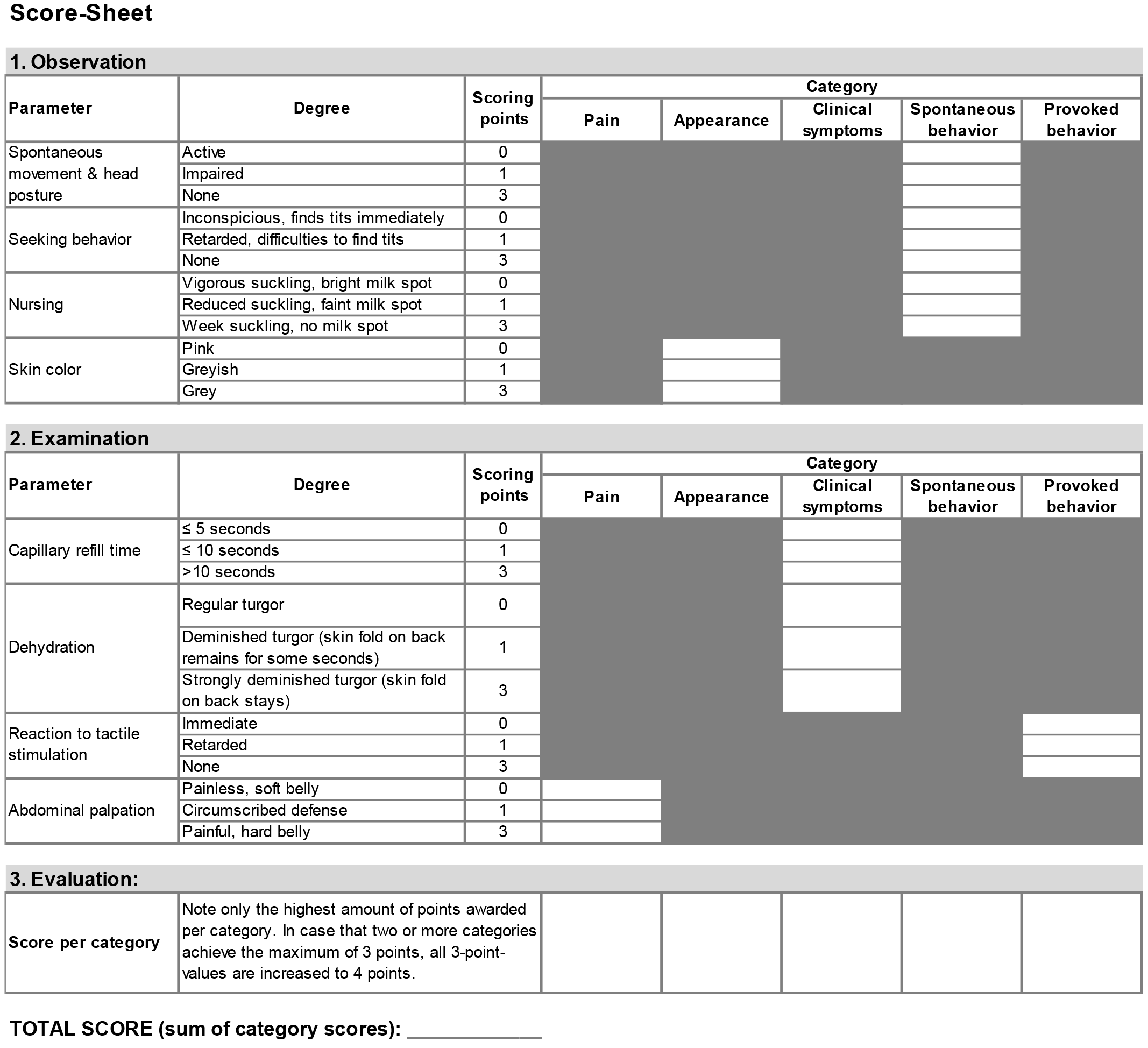
Score sheet. Documentation sheet for scores awarded to individual mice at relevant time points. Scoring requires observation and examination of indicated parameters to assess the different categorical aspects of the health status. The highest score values of each category are summed up to a final total score.

**FIG 2.**
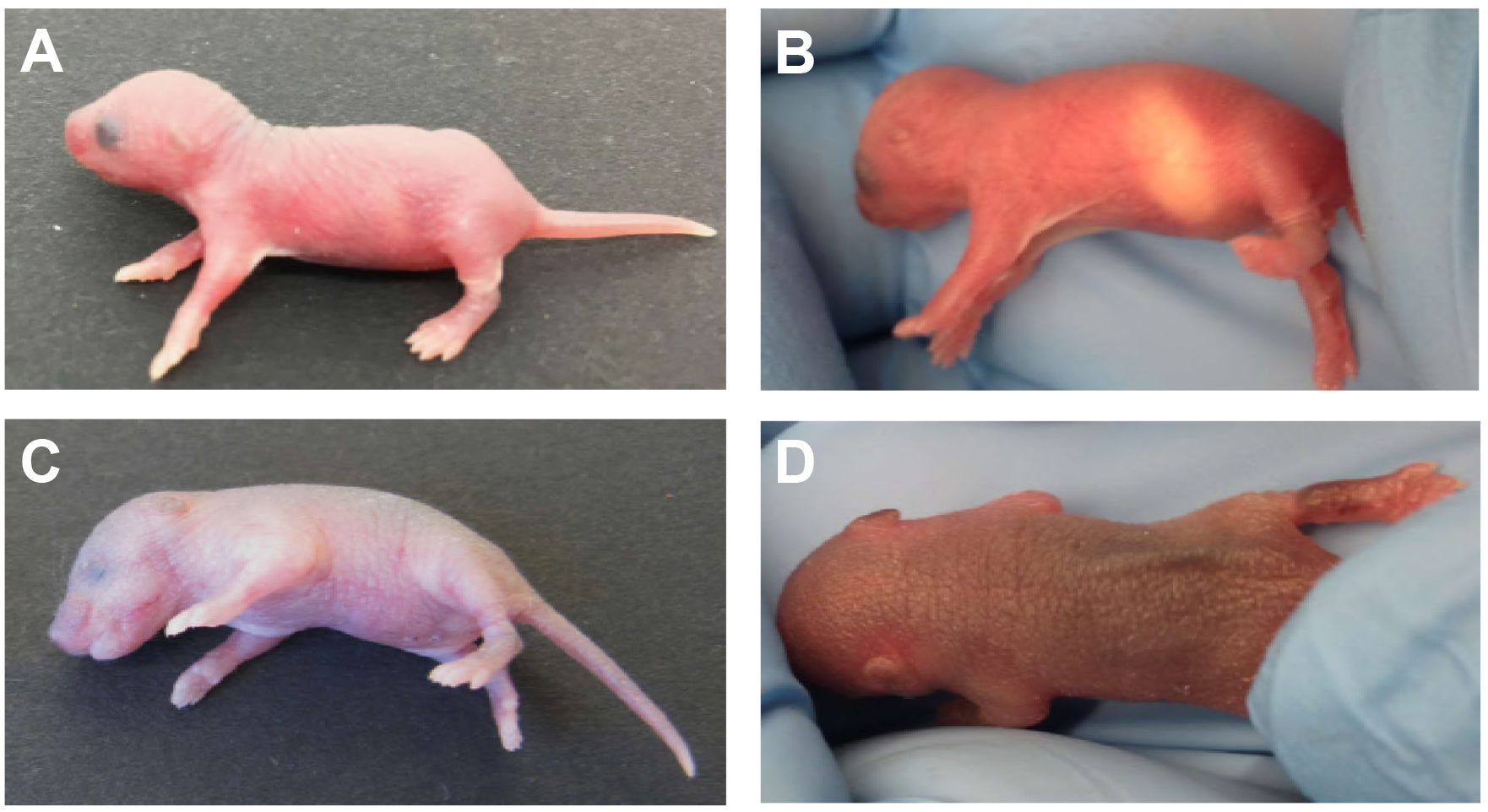
Typical appearance of observational scoring parameters in healthy and diseased neonatal mice. (A) Healthy newborn pup with active raised head posture, vivid spontaneous movement and rosy-pink skin color. (B) Bright abdominal milk spot indicating normal suckling behavior and nursing. (C) Neonatal mouse infected with *Staphylococcus (S.) aureus* presenting with impaired head posture, no spontaneous movement and greyish skin color. (D) Decreased skin turgor in a septic neonate with folds of the back skin remaining elevated due to dehydration.

### Clinical scoring correlates with sepsis-induced death rates in newborn mice

In order to validate the proposed scoring system in neonatal mice we applied an established model of *S. aureus*-induced neonatal sepsis (6). Two mouse strains (Wildtype (WT) and *s100a9*^*−/−*^) with different sepsis susceptibilities were used to test the reliability of the score. In this model, WT pups are significantly less prone to develop fatal sepsis than *s100a9*^*−/−*^ neonates (6). After challenge with *S. aureus*, mice were observed for 80 hours (h) and scored at defined times points. The distribution of awarded scores at the respective scoring time points is shown in Fig. 3A. During the first 12 h post infection (p.i.), most of the mice showed no significant symptoms (scores < 3). Subsequently, the proportion of scores > 3 slowly increased suggesting beginning of sepsis. Overt impairment of the health status resulted in score values of ≥5. Within the groups of evaluable, living mice at the respective time points, the proportion of scores ≥ 5 was over 50% from 28 h p.i. until 48 h p.i. The mean scores clearly correlated with the kinetics of death rates (spontaneous death and euthanasia) (Fig. 3B). These findings demonstrated that the scoring system was a reliable method to measure and to record the severity of the clinical course of sepsis in newborn mice.

**FIG 3.**
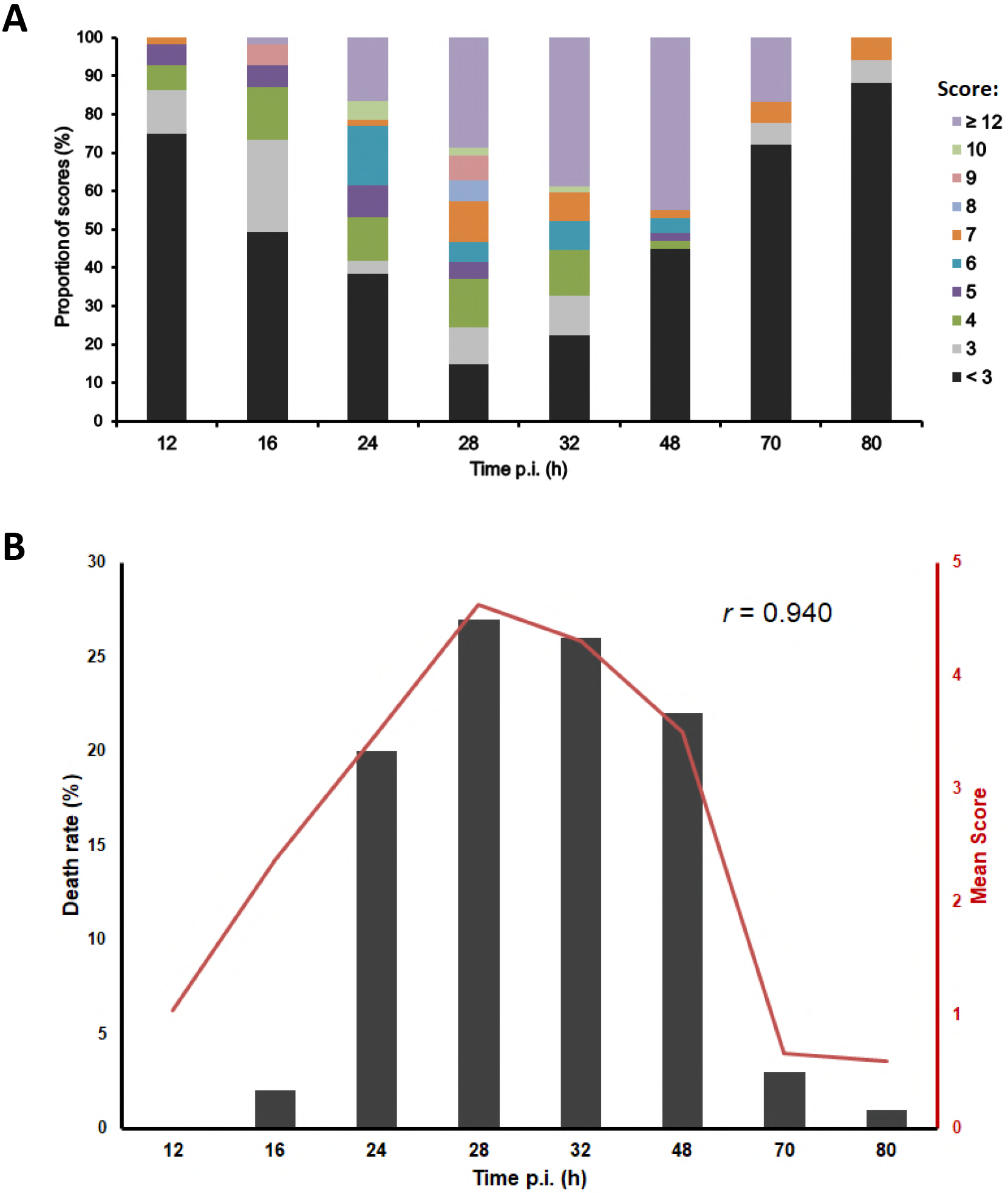
Clinical scoring results over the course of sepsis in neonatal mice. Sepsis was induced in neonatal mice (d2) (WT *n*=50, *s100a9^−/−^ n*=74) by subcutaneous injection of *S. aureus* (7 × 10^4^ CFU). (A) Bars show the distribution of awarded scores at the respective time points of examination. Scores are represented by distinct colors as indicated. (B) Bars show the death rates at indicated time points post infection (p.i.). The graphic line indicates the score mean at the respective time points of examination. *r*, Pearson’s correlation coefficient.

### The scoring system predicts fatal courses of sepsis in newborn mice

To assess the accuracy of the scoring system to predict fatal courses of sepsis we determined the highest score value each animal had been awarded during the observational period of 80 h. According to the highest score, animals were assigned to respective score maximum groups. Subsequently, the final outcome of the animals within these groups was determined and plotted as proportion of death and survival (Fig. 4). Overall this analysis revealed that 90% of mice with an at least once awarded score of ≥ 5 died (Fig. 4A). A score of ≥ 8 correlated with definitive fatality (100%). These results also hold true when looking separately at both of the mouse strains, i.e. the moderate sepsis susceptible WT neonates (Fig. 4B) and the highly sepsis susceptible *s100a9*^*−/−*^ neonates (Fig. 4C) (6). In neonates of both of these mouse strains, scores ≥ 5 indicated imminent deceasing and scores ≥ 8 unduly constituted fatal outcome. These findings demonstrated that the proposed clinical scoring system is a sensitive indicator of fatal courses of sepsis, allowing premature termination of experiments to prevent unnecessary suffer of neonatal experimental mice.

**FIG 4.**
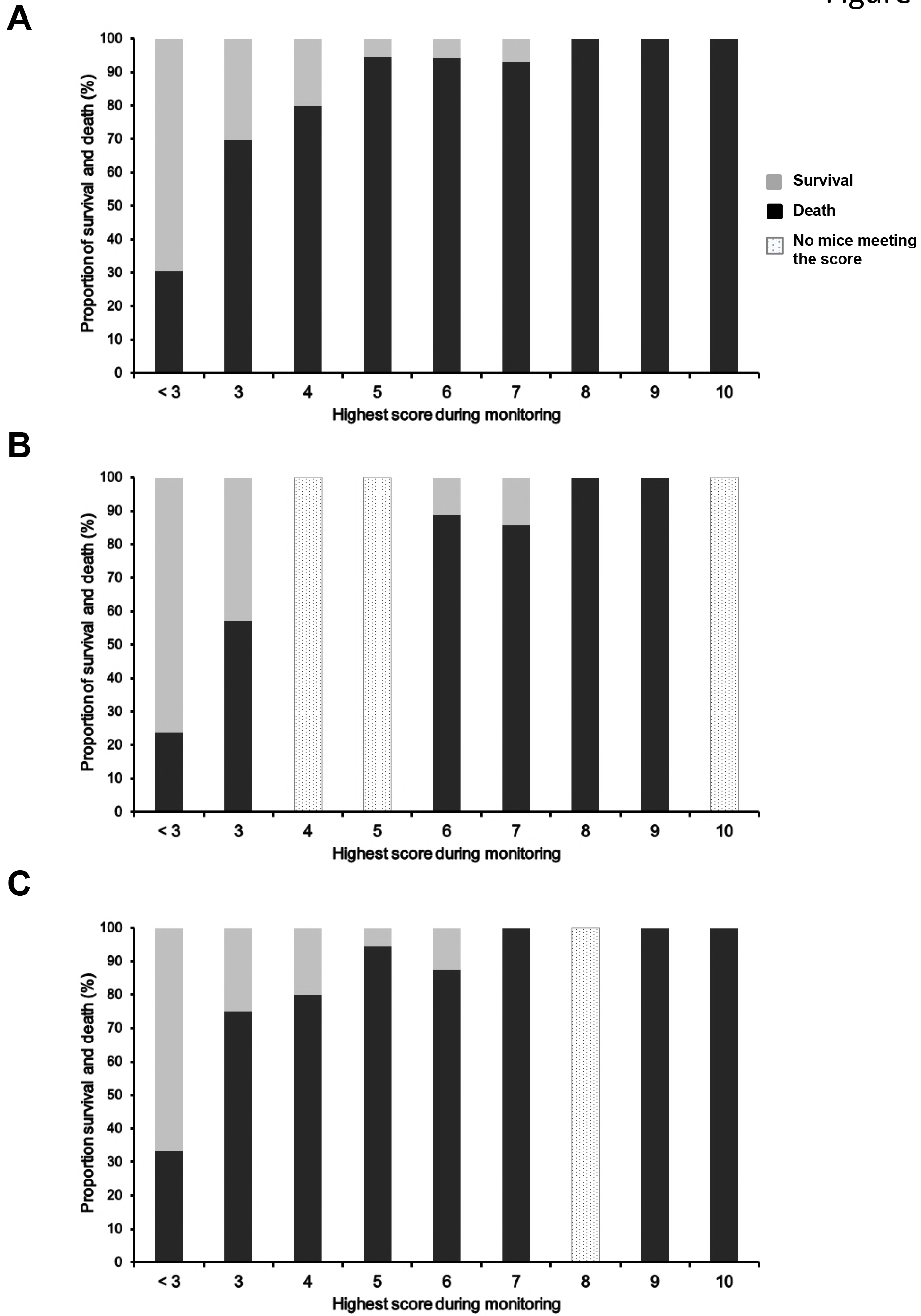
Clinical scoring predicts sepsis fatality in neonatal mice. After sepsis induction, the newborn mice were assigned to score maximum groups according to the highest score value they were awarded during the 80 hours of observation. Bars represent the proportion of survival and death within the score maximum groups. (A) Union of WT (*n* = 50) and *s100a9*^*−/−*^ (*n* = 74) neonates, (B) WT neonates only and (C) *s100a9*^*−/−*^ neonates only.

### Clinical scoring outperforms plasma cytokine levels in monitoring the disease state of septic newborn mice

In humans as well as in animal experiments, cytokine and chemokine levels in the plasma are used as biomarker for the severity of sepsis and the success of treatment (21–23). To corroborate the value of the proposed non-invasive scoring system we determined the plasma levels of Ccl7 (alias monocyte chemoattractant protein 3, Mcp-3), Ccl2 (alias Mcp-1), Il-6, Ccl5 (alias regulated on activation, normal T cell expressed and secreted, Rantes), Il-1α, and Tnf-α in *S. aureus*-infected neonatal mice 12 h p.i. (Fig. 5A) and 24 h p.i. (Fig. 5B). Cytokine levels were correlated with the scores awarded at these time points. At the early stage 12 h p.i., scores of *S. aureus*-infected mice were already increased (between 1 and 4) compared to the score mean + 2SD of PBS-treated control mice reflecting the beginning of sepsis. At this early time point, only the plasma levels of Ccl7 correlated well (*r* = 0.900) and those of Ccl2 mediocrely (*r* = 0.659) with the scoring results (Fig. 5A). The plasma levels of Il6, Ccl5, Il-1α and Tnf-α of infected mice did not correlate with the clinical scoring and were not increased compared to PBS-treated control mice (Fig. 5A). Only after 24 h of infection, the plasma levels of all cytokines correlated with the scoring results (Fig. 5B). However, in contrast to the clinical scoring, in most of the animals tested the levels of Ccl5, Il-1α and Tnf-α were not higher than in the control mice suggesting insufficient specificity for infection-induced increases (Fig. 5B). Collectively, the data demonstrated that clinical scoring of neonatal mice according to the proposed system is a highly sensitive early indicator of sepsis outmatching the value of common plasma cytokine levels as early biomarker. The correlation between scores and plasma cytokine levels at later stages of sepsis corroborates the quality of the scoring system as an objective follow-up parameter.

**FIG 5.**
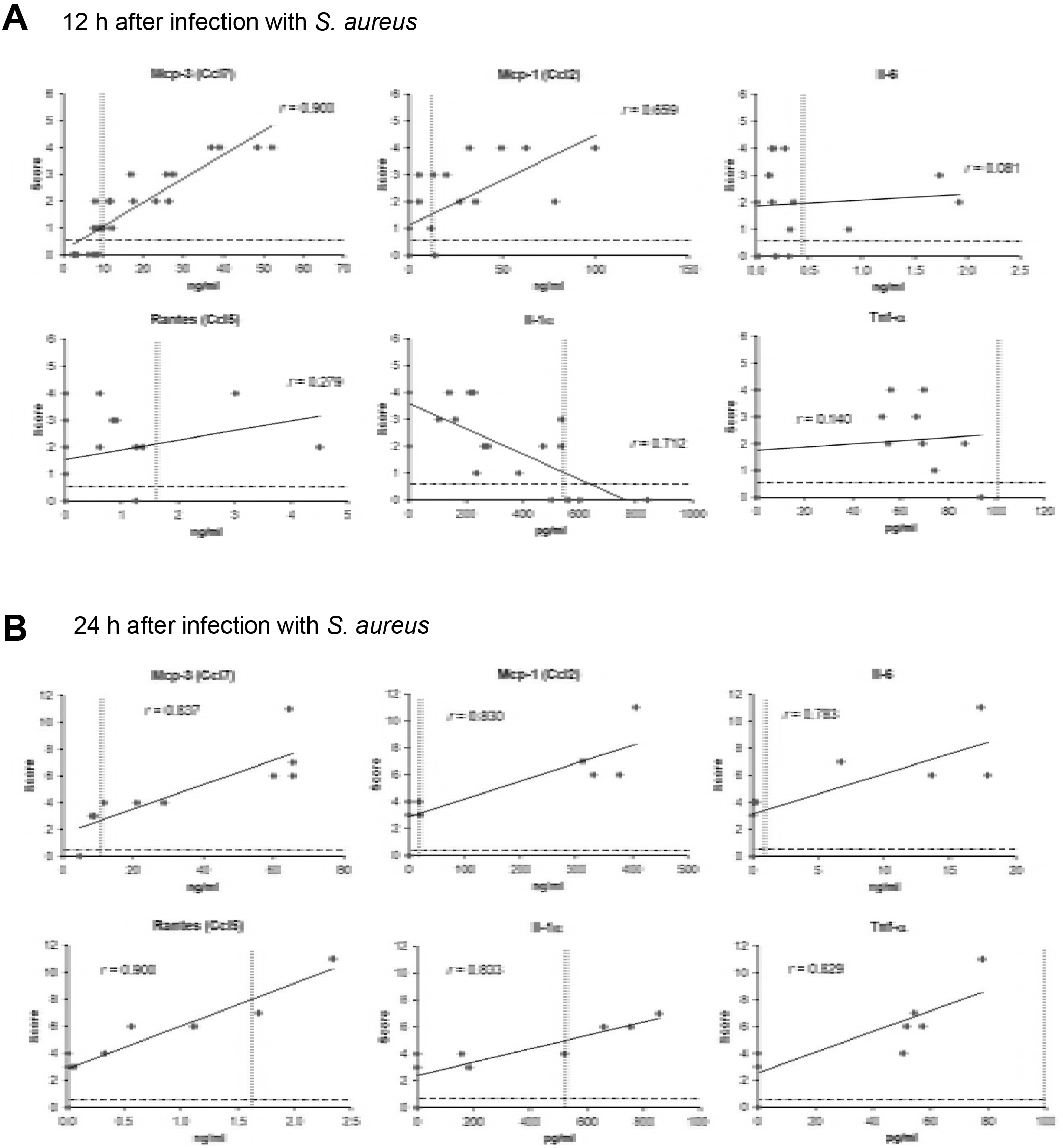
Clinical scoring is more sensitive than plasma cytokine levels to monitor early disease states in septic neonatal mice. Plasma cytokine levels of *S. aureus*-challenged WT (*n* = 19) and *s100a9*^*−/−*^ (*n* = 17) neonates (d2) were correlated with the scoring results in these mice as determined directly before killing at (A) 12 h and (B) 24 h after infection. Scatter plots indicate the best fit and the correlation coefficient *r*, respectively. Lines indicate the mean + 2SD of scores (dashed lines) and cytokine levels (dotted lines) in the group of PBS-injected control neonates.

## Discussion

Clinical scoring systems are widely used in animal research to evaluate the health status of experimental animals as an objective scientific parameter and for evaluating the animal welfare. In mice, scoring systems are only published for adults (15–17) but are missing in neonates. However, adult scoring systems are not readily transferable to neonatal mice since most of the clinical parameters used are not applicable in neonates due to absence of fur, posture, different movement patterns or the restricted accessibility of eyes. Even loss of weight is usually a useless parameter as their body weight increases physiologically in the first weeks of life in an exponential manner. Important scientific questions relating to early life immunity like the pathogenesis of neonatal sepsis and the imprinting impact of microbial challenges on the maturation of immunity (5–12) require the usage of animal models. Therefore, clinical scoring systems are needed that allow the reliable non-invasive evaluation of newborn experimental animals in a longitudinal manner.

In this study, we developed and validated a scoring system for newborn mice that closely followed clinical evaluation parameters used in human neonates. The clinical signs in diseased human neonates are usually unspecific and often subtle. The APGAR score is the most established score system in human neonates and includes <u>a</u>ppearance, heart rate (<u>p</u>ulse), responding to tactile stimuli (<u>g</u>rimace), <u>a</u>ctivity and <u>r</u>espiration (24). A more specific scoring system for septic human neonates is the Neonatal Therapeutic Intervention Scoring System (NTISS) that considers 62 parameters addressing to respiratory, cardiovascular and metabolic/nutritional impairments, extent of drug therapy and transfusions, need of monitoring and invasive procedures, and type and number of vascular accesses (25). Some characteristics of neonatal mice are of advantage; the absence of fur allows evaluating parameters inaccessible in adult mice, e.g. skin coloring or presence of milk spot as an indirect criterion for the suckling behavior. Based on these considerations, we developed the presented scoring system that demands observation and minimal physical examination of the newborn mice avoiding separation from the mother. Testing the proposed scoring system in two different mouse strains that underwent neonatal sepsis induced by *S. aureus* showed a highly significant correlation of the disease course and death kinetics with increasing score values. Moreover, clinical scoring was able to predict sepsis fatality. The analysis of the outcome of individual neonatal mice revealed a 90% death rate if a score value of ≥ 5 had been achieved. Mice which were evaluated with a score of ≥ 8 died at 100%. Finally, the correlation of the scores with plasma cytokine levels in *S. aureus*-infected and PBS-treated control mice demonstrated that plasma cytokine levels are less sensitive than non-invasive scoring to indicate beginning sepsis in neonates. After 12 h of treatment, the plasma levels of Il-6, Ccl5, Il-1α and Tnf-α of infected and control neonates did not differ whereas the scoring values of the infected pups were already increased. Only after 24 h of sepsis induction all plasma cytokine levels correlated with the scoring results. Therefore, at an early stage, clinical scoring is more helpful than the analysis of plasma cytokine levels to detect beginning sepsis in neonatal mice. These findings were in line with the well-known dilemma in human neonates that cytokine levels insufficiently indicate imminent septic events while clinical evaluation is still the most sensitive method (18, 19, 26). Of note Il-6, the most common sepsis biomarker used in human neonates, was only at the late stage of sepsis (24 h p.i.) increased in neonatal mice.

In order to ensure wellbeing and protection of experimental animals monitoring schedules with meaningful time points of assessment have to be established. Initially, it is therefore mandatory to determine when first disease signs in the respective disease model occur and death rates peak. The clinical scoring system proved as a valuable and highly sensitive tool detecting impairments of the health status at the earliest conceivable moment. In the sepsis model used in this study, death rates started to peak after 24 h p.i. and remained high until 48 h p.i. (20-27%). For this model, we therefore propose a scoring schedule (Table 1) that includes close monitoring during the first two hours after starting the experiment to detect unforeseeable reactions and experimenter errors like unintended intravenous injection or incorrect bacterial dosage. This is followed by a period when scoring every 4 hours is sufficient. After 20 h p.i., monitoring should be intensified again to every 2 hours to detect unnecessary suffering from sepsis by predicting fatality based on the scoring result that demands euthanasia. Animals surviving 48 h p.i. are unlikely to die anymore from infection, wherefore further monitoring three times a day is sufficient until end of experiments.

**Table 1.**
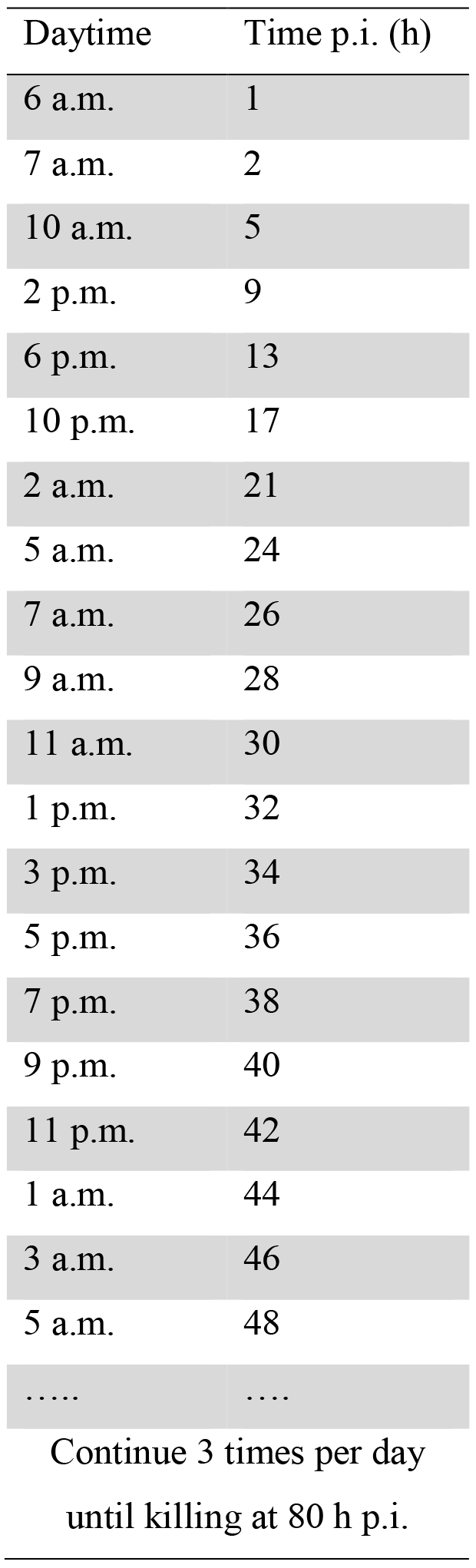
Monitoring schedule of neonatal mice after sepsis induction with *Staphylococcus aureus*.

In summary, the proposed scoring system is the first reliable and sensitive tool that detects health impairments in septic neonatal mice at an early disease stage and predicts fatal outcomes based on clinical observation and minimal physical examination. In newborn mice, clinical scoring according to the here validated system allows longitudinal assessment of the health status and is ahead of common invasive laboratory markers of inflammation to detect imminent sepsis.

## Materials and Methods

### Mice

C57BL/6 mice (wildtype, WT) and *s100a9* knock-out mice (−/−) (27) were used for breeding and housed under specific pathogen-free conditions in the same animal room at the University of Veterinary Medicine Hannover, Germany and maintained under standard conditions according to institutional guidelines. The *s100a9*^*−/−*^ mouse is a widely-accepted C57BL/6 mouse strain (27) and meanwhile these mice have been backcrossed to F12 generation of C57BL/6 background (5, 28, 29). Both mouse strains were constantly bred, and litters were used randomly. For experiments, neonates were used at the age of 2 days (d2) and only if the mothers had already given birth three times at minimum. No animals needed to be excluded from the studies.

### Ethical approval

Mouse experiments were carried out in accordance with German Animal Welfare Legislation and performed as approved by the Lower Saxony State Office for Consumer and Food Safety, Germany (approval no. 33.12-42502-04-14 and no. 33.12-42502-04-15).

### Sepsis model

Neonatal d2 WT and *s100a9*^*−/−*^ mice were carefully lifted from their nest and placed onto soft papers on a styrofoam underground. Forceps with rubber tips were used to hold the animal tight for treatment. Sepsis was induced by subcutaneous injection of 20 μl of bacterial suspensions containing 7 × 10^4^ CFU *S. aureus* strain Newman (GenBank accession number AP009351.1) in the back of the neonates. Mice injected with 20 μl PBS served as controls. The procedure was performed calm and uninterrupted within minutes to ensure reacceptance of the neonate by the mother. Mice were monitored for survival over a time period of 80 h at maximum. For cytokine studies, mice were sacrificed by decapitation 12 h and 24 h after bacterial inoculation to harvest blood. Monitoring and scoring was performed every hour during the first 12 h p.i. and then every 2 h until killing.

### Scoring system for neonatal mice

Animals were observed for at least 5 min to assess spontaneous movement and head posture, seeking behavior, nursing and skin color. Parameters that required physical examination were the capillary refill time to evaluate the circulatory status, the skin turgor to detect signs of dehydration, tactile stimulation to test flight reflexes and alertness, and abdominal palpation to assess unconscious defense reactions due to pain. Appraisal of the milk spot shining from the stomach through the abdomen (30) was also considered when analyzing the nursing behavior. For each parameter scoring points were awarded for deficiencies of the health status, rated in terms of no (0 point), moderate (1 point) or significant (3 points) impairment. The highest value of scoring points awarded per category was noted and added up to the final score. If the maximum of 3 points had been awarded twice or more, all 3-point-values were increased to 4 points. Animals were euthanized by painless decapitation if the final score reached at least 12 points or if a strong drinking deficiency with no seeking behavior for more than 12 hours was observed.

### Cytokine assays

Blood was collected using heparinized glass capillaries and transferred into heparinized tubes. After centrifugation at 500 × g for 5 min, plasma was removed and centrifuged at 2000 × g for 5 min and stored at −80 °C until cytokine analysis was performed using the LEGENDplex Mouse Multi-Analyte Flow from BioLegend (San Diego, USA) according to manufactures’ instructions. Samples were analyzed with a FACS Canto II flow cytometer (BD Biosciences). Data were processed using DIVA software v8.0.1 (BD Biosciences) and LEGENDplex Data Analysis Software v7.0 (BioLegend).

### Statistics

For correlation analyses, linear regression was computed, Pearson’s correlation coefficient *r* determined and trend lines (best fits) indicated.

## Acknowledgement

We thank Sabine Schreek for excellent technical support. This work was supported by grants from the Appenrodt Foundation, the German Research Foundation [VI 538/6-1]; and the Volkswagen Foundation [Az 90005] to DV. The funders had no role in the study design, data collection, interpretation and analysis, decision to publish, or preparation of the manuscript.

